# A systematic analysis of hypermucoviscosity and capsule reveals distinct and overlapping genes that impact *Klebsiella pneumoniae* fitness

**DOI:** 10.1101/2020.06.22.164582

**Authors:** Laura A. Mike, Andrew J. Stark, Valerie S. Forsyth, Jay Vornhagen, Sara N. Smith, Michael A. Bachman, Harry L.T. Mobley

**Author notes:** **Corresponding authors**, (LAM) and (HLTM). These authors contributed equally to this work (Andrew J. Stark, Valerie S. Forsyth).

## Abstract

Hypervirulent *K. pneumoniae* (hvKp) is a distinct pathotype that causes invasive community-acquired infections in healthy individuals. Hypermucoviscosity (hmv) is a major phenotype associated with hvKp characterized by copious capsule production and poor sedimentation. Dissecting the individual functions of CPS production and hmv in hvKp has been stymied by the conflation of these two properties. Although hmv requires capsular polysaccharide (CPS) biosynthesis, other cellular factors may also be required and some fitness phenotypes ascribed to CPS may be distinctly attributed to hmv. To address this challenge, we systematically identified genes that impact capsule and hmv. We generated a condensed, ordered transposon library in hypervirulent strain KPPR1, then evaluated the CPS production and hmv phenotypes of the 3,733 transposon mutants, representing 72% of all open reading frames in the genome. We employed forward and reverse genetic screens to evaluate effects of novel and known genes on CPS biosynthesis and hmv. These screens expand our understanding of core genes that coordinate CPS biosynthesis and hmv, as well as identify central metabolism genes that distinctly impact CPS biosynthesis or hmv, specifically those related to purine metabolism, pyruvate metabolism and the TCA cycle. Six representative mutants, with varying levels of CPS production and hmv, were all significantly out-competed by wildtype in a murine model of disseminating pneumonia. This suggests that an optimal balance between cellular energetics, CPS biosynthesis and hmv are required for maximal fitness. Altogether, these data demonstrate that hmv requires both CPS biosynthesis and other cellular factors, and that these processes are integrated into the metabolic status of the cell. Therefore, hvKp may require certain nutrients to fully elaborate its virulence-associated properties to specifically cause deep tissue infections.

**Author summary:** *Klebsiella pneumoniae* is a common multi-drug resistant hospital-associated pathogen, however some isolates are capable of causing community-acquired infections in otherwise healthy individuals. The strains causing community-acquired infections have some distinguishing characteristics, which include overproduction of capsule and hypermucoviscosity. Hypermucoviscous strains are very tacky and sediment poorly when centrifuged. Historically, hypermucoviscosity has been attributed to overproduction of capsular polysaccharide, but recent data suggest that other factors contribute to this bacterial phenotype. Moreover, it seems that capsule and hypermucoviscosity may have distinct roles in pathogenesis. In this study, we sought to systematically investigate the genes that contribute to capsule and hypermucoviscosity. We found that in most cases, genes coordinately impact both capsule biosynthesis and hypermucoviscosity. Some metabolic genes linked to the TCA cycle, however, only affect one of these properties. Here, we identify that capsule biosynthesis and hypermucoviscosity are tightly tied to central metabolism and that an optimal balance between metabolism, capsule, and hypermucoviscosity are important for *in vivo* fitness of *K. pneumoniae*. These results identify genes that can be further probed to dissect how capsule and hypermucoviscosity are coordinated in response to niche-specific nutrients. Such studies will expand our understanding of the factors that drive the pathobiology of hypervirulent *K. pneumoniae*.

## Introduction

*Klebsiella pneumoniae* is a ubiquitous bacterium found in a range of environments, including soil, sewage, sink P-traps, and mammalian gastrointestinal tracts. Colonization of the human gut with *K. pneumoniae* is a risk factor for infection, which commonly manifests as hospital-associated pneumonia, urinary tract infections, and bacteremia [1–3]. Classical *K. pneumoniae* (cKp) is commonly an opportunistic pathogen causing infections in patients who are immunocompromised, have indwelling medical devices, have undergone an invasive medical procedure, or have other co-morbidities such as diabetes mellitus and alcoholism [4, 5]. With human colonization rates reported at 23-36%, increasing antibiotic resistance, and a non-fastidious lifestyle, it is not surprising that *K. pneumoniae* is the third most common nosocomial pathogen [1, 3, 4, 6].

Two clinically challenging pathotypes with high morbidity and mortality are the carbapenem-resistant, classical *K. pneumoniae* (CR-cKp) and hypervirulent *K. pneumoniae* (hvKp) [5, 7–9]. CR-cKp was first observed in 1996 and since then has been the major driving force disseminating carbapenem-resistance throughout the Enterobacteriaceae, complicating the treatment of many gram-negative infections [10, 11]. In parallel, hvKp incidence is rising in both community and hospital settings [12–15]. While hvKp is susceptible to most antibiotics, it is associated with invasive infections in otherwise healthy patients and is notorious for causing pyogenic liver abscesses and disseminating to the eyes, lungs and brain, a pathogenesis uncommon for gram-negative enteric bacteria [3, 7, 8]. HvKp mortality rates range from 3 to 55% and survivors often have severe morbidities such as vision loss or neurologic sequelae [7, 8]. Alarmingly, the CR-cKp and hvKp pathotypes can converge [3, 16]. The prevalence of CR-hvKp is 7.4-15% in countries where hvKp is endemic, demonstrating that more devastating *K. pneumoniae* lineages are emerging [7].

Accessory features associated with hvKp include hypermucoviscosity (hmv), K1 or K2 capsule-types, overexpression of RmpA (regulator of mucoid phenotype), and stealth siderophore biosynthesis [7, 12, 14, 15, 17, 18]. Traditionally, *K. pneumoniae* isolates are categorized as hmv by string test if their colony stretches more than five mm when picked off a plate (**Fig 1B**). In addition, overexpression of RmpA has been shown to increase capsular polysaccharide (CPS) production [19, 20]. A clear link between CPS and virulence has been demonstrated in multiple murine models of *K. pneumoniae* infection, including pneumonia and UTI [21, 22]. Despite CPS being a key fitness factor for *K. pneumoniae*, the regulatory network that directly controls CPS biosynthesis is not fully understood. A recent study reported the *K. pneumoniae* CPS biosynthesis regulatory network using density-TraDISort [23]. Transposon mutant pools were separated over a discontinuous Percoll gradient to separate populations with altered buoyancy as a surrogate measure of CPS production in two hvKp strains, NTUH-K2044 and ATCC 43816 [23]. Transposon insertions were identified that increase buoyancy in NTUH-K2044 or decrease buoyancy in NTUH-K2044 and/or ATCC 43816, then the hmv and CPS production of ten targeted deletion mutants were quantified to validate the density-TraDISort. Building upon this study, we sought to systematically expand our understanding of the relationship between CPS biosynthesis and hmv using all available genes of interest identified by density-TraDISort.

**Fig 1.**
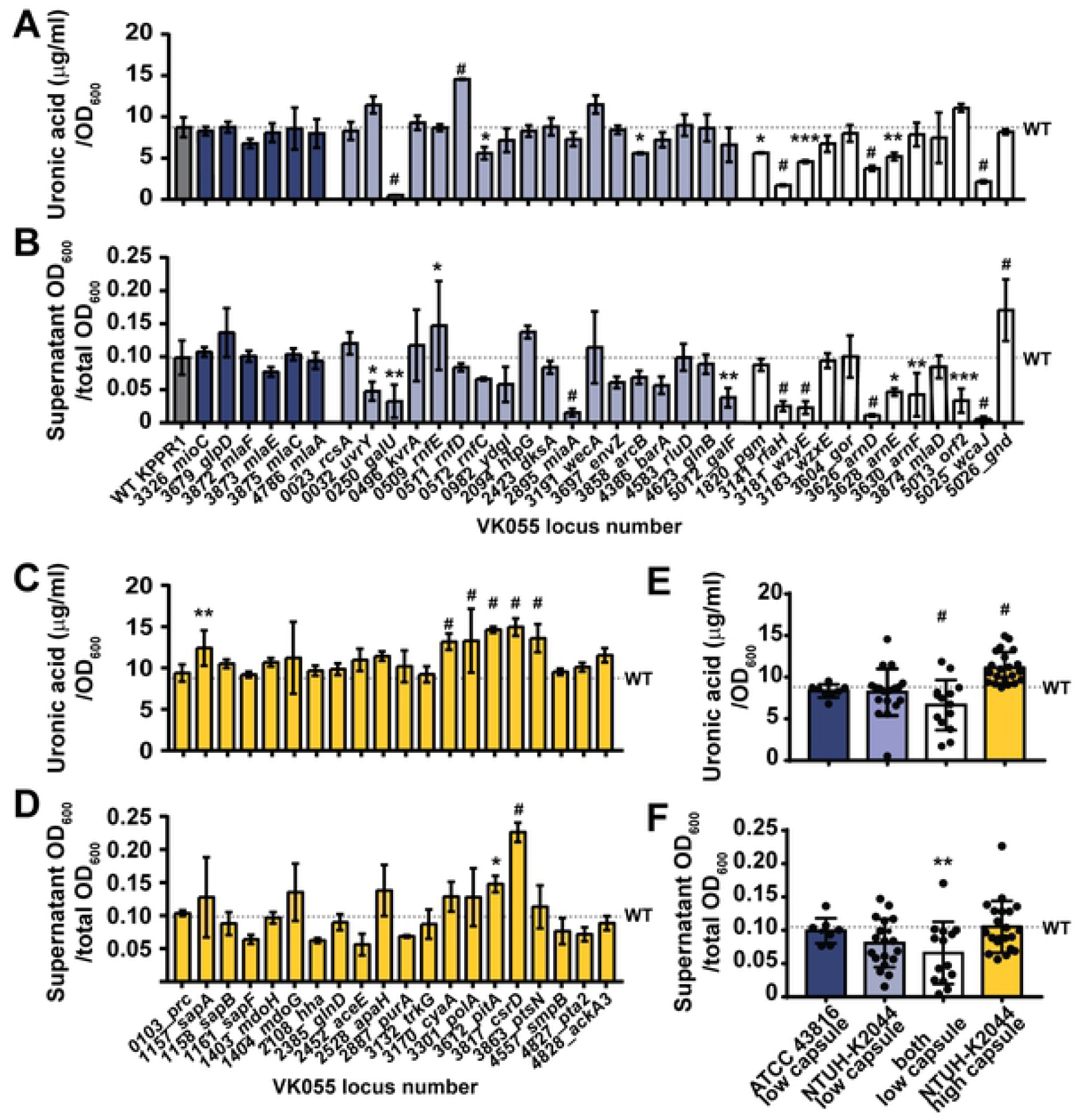
A forward phenotypic screen of the condensed, ordered *K. pneumoniae* library. (A) A Mariner *Himar1* transposon (Tn) library was generated in *K. pneumoniae* strain KPPR1 and arrayed into 192, 96 well microplates. 14,895 traceable Tn insertions were mapped, disrupting 71.6% of open reading frames (ORFs) and 74.2% of predicted transcriptional units. The best representative Tn insertion for the 3,733 disrupted genes were consolidated into a condensed library. 58% of all Tn insertion sites are located within the first 66.7% of each ORF. (B) The condensed transposon library was then screened for non-mucoid mutants as indicated by improved sedimentation. Strains with supernatant OD_600_ values two standard deviations below the plate mean were then evaluated on solid medium by string test. Each hit was arrayed in triplicate into 96 well plates then confirmed by sedimentation and string test.

Historically, hmv has been closely associated with hvKp and attributed to over-production of CPS, as hmv is lost when CPS biosynthesis is reduced [7]. This paradigm is pervasive throughout the *K. pneumoniae* literature despite an early study suggesting that hmv may not only be due to overproduction of CPS [24]. More recently, discordant changes in CPS production and hmv have been shown at the phenotypic and genotypic levels [3, 25]. Some examples include the *rmpC* mutant in strain KPPR1S and the MagA_G334A_ (<u>m</u>ucoviscosity-<u>a</u>ssociated <u>g</u>ene <u>A</u>, also known as Wzy) mutant in strain NTUH-K2044, which both exhibit reduced CPS production, but remain hmv [26–29]; and the *rmpD* mutant in strain KPPR1S and the *kvrA* (*<u>K</u>lebsiella* <u>v</u>irulence <u>r</u>egulator) mutant in the ST258 strain MKP103, which remain fully encapsulated, but are non-mucoid [25, 30]. The mounting evidence that hmv and CPS overproduction has been conflated into a single characteristic of hvKp spotlights our limited understanding of *K. pneumoniae* hypervirulence and points to the critical need to better understand the relationship and function of these two important features in *K. pneumoniae* pathogenesis and biology.

To systematically evaluate the relationship between CPS biosynthesis and hmv and provide a robust resource for future molecular studies, we have developed an ordered transposon library in the hypervirulent *K. pneumoniae* strain KPPR1 using Cartesian-Pooling and Coordinate-Sequencing [31]. We then condensed the library to include a representative mutant for each of the 3,733 disrupted genes. To validate the use of the library and more broadly examine the relationship between hmv and CPS overproduction, forward and reverse genetic screens were performed to systematically quantify both CPS production and hmv exhibited by transposon mutants. The use of a forward screen allowed for the unbiased identification of mutants that impact hmv and/or CPS production, while the reverse screen enabled methodical screening of CPS production and hmv for genes previously ascribed to affect *K. pneumoniae* buoyancy [23]. Global analyses of the CPS production and hmv of 100 transposon mutants and 27 targeted mutants revealed a significant correlation between the two biological features, although several mutants displayed discordant regulation of hmv and CPS biosynthesis. These data strengthen the emerging model that CPS production and hmv are tightly linked, but distinct; emphasizing the need to decouple these features and define their individual contributions to hypervirulence. Since hvKp typically cause invasive infections that exhibit metastatic spread [5], we used a murine model of disseminating pneumonia to begin addressing this question. We evaluated the *in vivo* fitness of six representative mutants with an array of hmv and CPS levels in this model. All mutants were significantly out-competed *in vivo*, suggesting that coordinated regulation of CPS biosynthesis and hmv are critical for maximal virulence. Therefore, it is of utmost importance that the overlapping and distinct pathways controlling hmv and CPS biosynthesis be further mapped so that the functional relationships between hmv, CPS, and hypervirulence in *K. pneumoniae* can be further dissected. Such studies may ultimately identify anti-virulence targets useful for specifically treating hvKp infections

## Results

### Generating a condensed, ordered transposon library in *Klebsiella pneumoniae* strain KPPR1

To facilitate both forward and reverse genetic studies in *K. pneumoniae*, an ordered transposon library of strain KPPR1, a rifampin-resistant derivative of ATCC 43816 that has a K2 capsule type and exhibits hypermucoviscous behavior, was generated [32]. Mariner *Himar1* transposon mutants were arrayed into 192, 96-well microplates and the transposon insertion site present in each well of the library was identified using Cartesian Pooling and Coordinate Sequencing (CP-CSeq) [31]. The library contains 14,895 traceable transposon mutants that disrupt 3,733 genes, covering 71.6% and 74.2% of predicted open reading frames (ORFs) and transcriptional units, respectively (**Table 1** and **Supplemental Data Set 1**). For each gene disrupted, a representative transposon mutant was selected to generate a condensed library (**Supplemental Data Set 1**). The representative mutant for each ORF was selected based on the confidence of its unique positional location in the library and the proximity of the transposon insertion site to the translational start site; 58% of all transposon insertion sites are in the first 66.7% of the ORF (**Fig 1A**). The selected mutants, representing 87% of KPPR1 non-essential genes, were arrayed into 41 microplates (**Fig 1A** and **Supplemental Data Set 1**) [33]. The accuracy of CP-CSeq identification of mutant positional locations was evaluated by PCR, where 92.9% (N = 14) and 93.8% (N = 16) of tested transposon mutants from the complete and condensed libraries, respectively, were confirmed to have the expected transposon insertion site. It is important to recognize that (1) absence of PCR product does not preclude the possibility that the correct transposon insertion site was present, but not detected by the PCR, (2) presence of PCR product does not exclude the possibility of additional transposon insertion sites sharing the library location, and (3) one mutant that did not validate in the condensed library grew poorly.

**Table 1.**
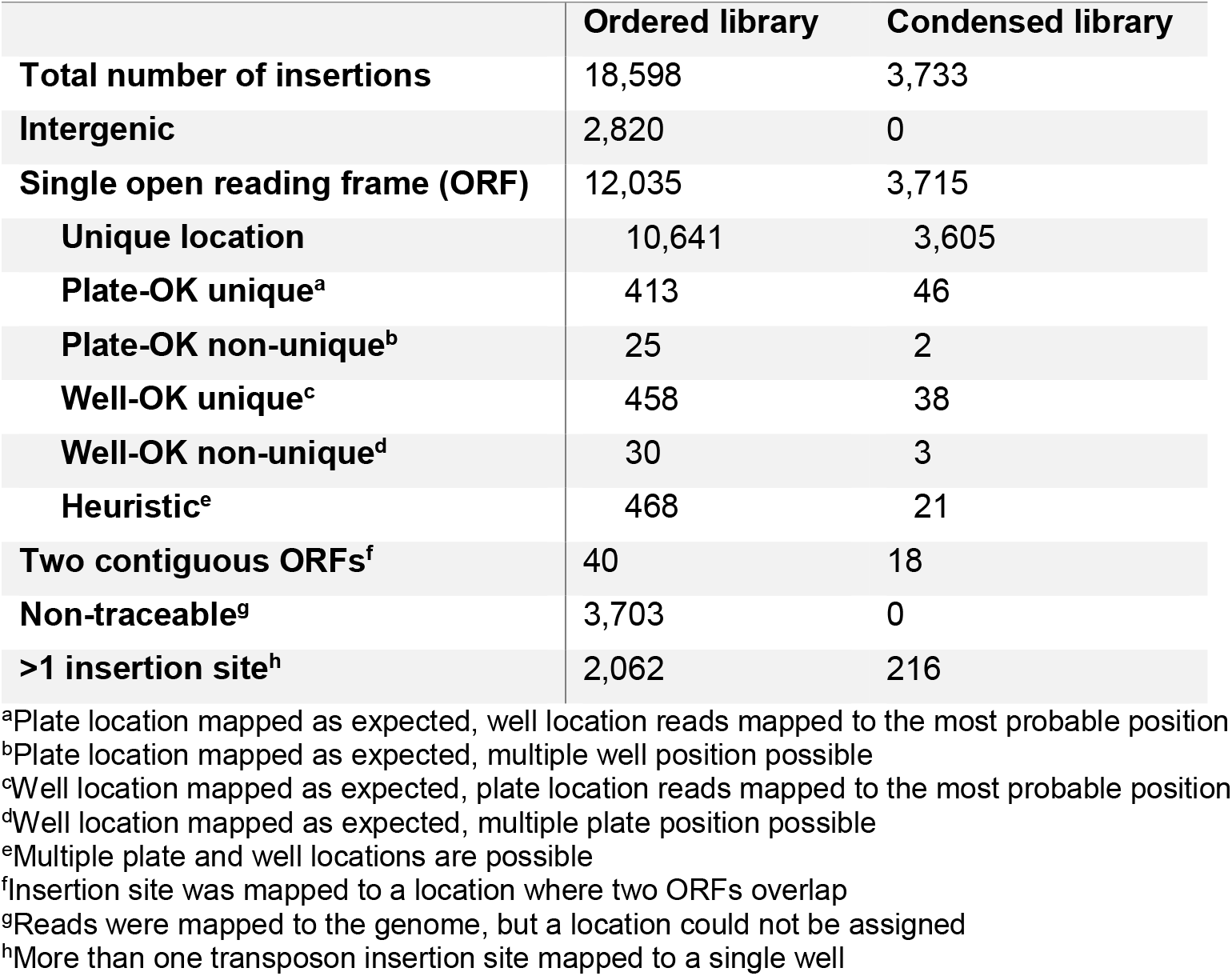
Composition of ordered and condensed *K. pneumoniae* strain KPPR1 transposon libraries.

### An unbiased forward phenotypic screen identified genes that influence hypermucoviscosity and capsular polysaccharide production

The classification of an isolate as hmv is typically done by a string test, where a colony is lifted off a plate with an inoculating loop and if it stretches more than five mm it is considered hmv (**Fig 1B**). *K. pneumoniae* hmv can also be quantified in liquid cultures by sedimentation since hypermucoviscous cells are retained in the supernatant after centrifugation, while non-mucoid cells fully sediment to form a tight pellet [25, 34]. This objective assay is more quantitative and reproducible than the string test.

To validate the utility of the transposon library by identifying both known and novel genes that impact *K. pneumoniae* hmv, the condensed library was screened for transposon mutants with reduced hmv using sedimentation assays and the string test (**Fig 1B**). With a hit rate of 2.76%, the 103 mutants initially identified to have reduced hmv based on sedimentation were patched onto LB plates. There, loss of hmv was confirmed in a secondary screen by string test followed by a sedimentation assay performed in triplicate (**Fig 1B**). 53 of the 103 primary hits passed these confirmation assays and were then evaluated in a final sedimentation assay scaled up to a standard culture volume of 3 mL (**Fig S1**). Ultimately, 44 hypo-mucoid (hmv^low^) transposon mutants passed the 3 rounds of screening and confirmation (**Table 2**). Most of the hits did not stretch at all by string test and were therefore sub-categorized as non-mucoid (hmv^0^, 33 hits), leaving 11 hmv^low^ hits that stretched less than five mm by string test (**Fig S1** and **Table 2**).

**Table 2.**
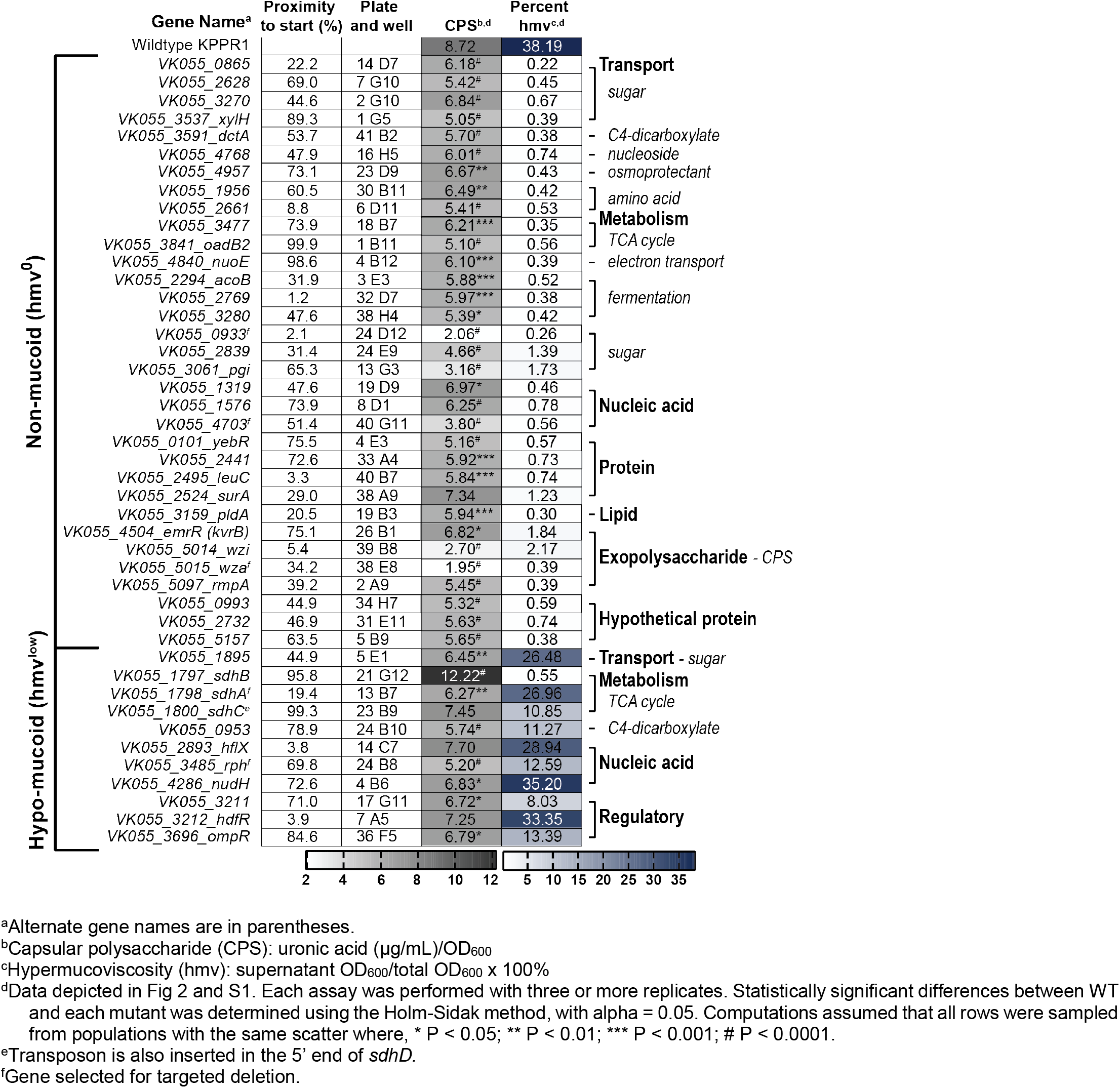
Forward phenotypic screen of KPPR1 hypermucoviscosity and capsular polysaccharide biosynthesis.

To investigate the relationship between hmv and CPS production, all 44 transposon mutants were evaluated for capsule (CPS) production by quantifying the amount of uronic acid produced by each strain (**Fig 2**) [35]. Overall, both classes of transposon mutants produced significantly less uronic acid than WT, although hmv^0^ hits produced significantly less CPS than hmv^low^ hits (**Fig 2A**). Specifically, 97.0% of the hmv^0^ (N = 33) and 63.6% of the hmv^low^ (N = 11) hits synthesized significantly less CPS than WT (**Fig 2**). Notably, hmv^0^ hits encompassed a wide range of CPS levels, yet were all non-mucoid, and many hmv^low^ strains produced quantities of CPS comparable to hmv^0^ strains, yet retained some mucoidy (**Fig 2** and **S1**). Altogether, these data support that CPS production is necessary for *K. pneumoniae* to exhibit hypermucoviscosity, yet emphasize that other bacterial factors are also likely required for hmv (**Fig 2A**).

**Fig 2.**
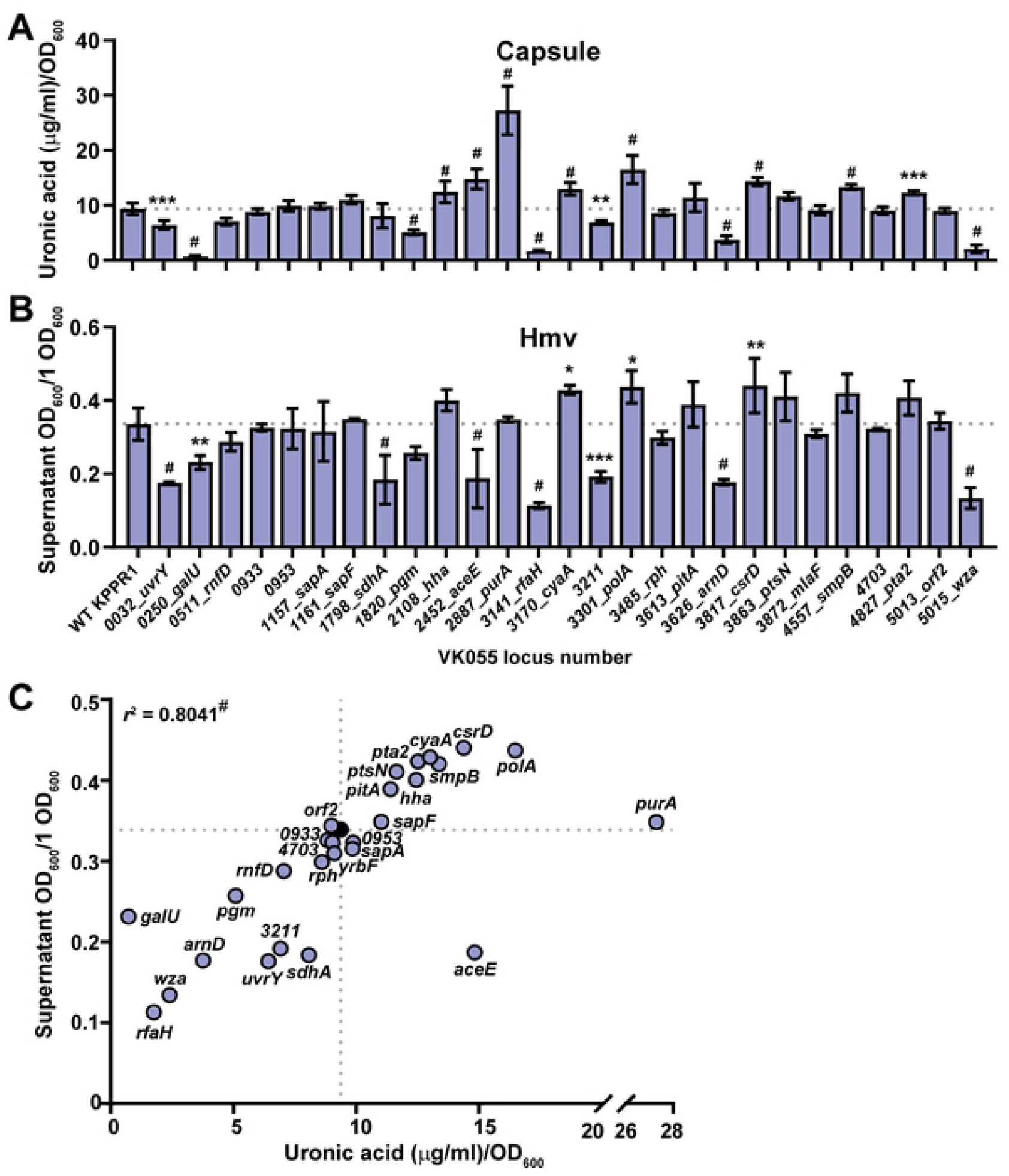
A forward screen identifies transposon mutants that influence *K. pneumoniae* hypermucoviscosity (hmv) and capsular polysaccharide production. (A) The mean capsular polysaccharide (CPS) production of 11 hypo-mucoid (hmv^low^) and 33 non-mucoid (hmv^0^) transposon mutants were quantified in triplicate by measuring uronic acid content and normalized to the optical density at 600 nm (OD600). Each marker for wildtype (WT) is an individual replicate (N = 24). Error bars represent one standard deviation from the mean and statistical differences between WT and classes of mutants were determined by unpaired *t*-test. The mean uronic acid content of each (B) hmv^low^ and (C) hmv^0^ transposon mutant represented by a dot in (A) is shown individually and labeled with the old locus tag number and gene name. Error bars represent one standard deviation from the mean of the assay performed in triplicate. Statistical significance between WT (gray bar or dotted line) and each mutant was determined using the Holm-Sidak method, with alpha = 0.05. Computations assumed that all rows were sampled from populations with the same scatter where, * P < 0.05; ** P < 0.01; *** P < 0.001; # P < 0.0001. The mean WT value is denoted by a horizontal, dotted gray line.

The forward transposon screen identified six genes previously identified to support CPS production, including five hmv^0^ hits (*wzi, wza, rmpA, kvrB*, and *pgi*) and one hmv^low^ hit (*ompR*) (**Fig 2C**) [23, 24, 30]. These findings serve as an internal experimental validation, which provide confidence in our results obtained from screening the transposon library. In addition, the unbiased forward screen identified other genes involved in central metabolism and bacterial cell biology that have not been previously ascribed to impact CPS biosynthesis and hmv. The two classes of genes with the most hits included those involved in cellular metabolism (N = 13) and transport (N = 10), half of which are predicted to have cognate sugar substrates (**Fig 2B-C, Table 2**). The ten transporters identified include five sugar transporters (*VK055_0865, VK055_2628, VK055_3270, xylH, VK055_1895*), a C4-dicarboxylate transporter (*dctA*), two amino acid transporters (*VK055_1956, VK055_2661*), a nucleoside transporter (*VK055_4768*), and an osmoprotectant transporter (*VK055_4957*) [36, 37]. Furthermore, 13 genes that participate in central metabolism were hit including those involved in TCA cycle (*VK055_3477, oadB2, sdhB, sdhA, sdhC*), electron transport (*nuoE*), fermentation (*acoB, VK055_2769, VK055_3280*), C4-dicarboxylate metabolism (*VK055_0953*), and sugar metabolism (*VK055_0993, VK055_2839, pgi*), along with many of the aforementioned genes involved in the transport of substrates for these metabolic processes [36, 37]. Intriguingly, eight genes related to nucleic acid function were identified including DNA replication, transcription, and RNA biology (*VK055_1319, VK055_1576, hflX, VK055_4703, VK055_3211, hdfR, rph*, and *nudH*), as well as four genes related to protein biology (*yebR, VK055_2441, leuC*, and *surA*) and one related to lipid biology (*pldA*) [36, 37]. Three hypothetical genes were identified, including *VK055_0933, VK055_2732*, and *VK055_5157*. Note that the apparent increase in CPS production in the *sdhB* transposon mutant may be confounded by an overt growth defect which dramatically skewed the normalization of uronic acid production to OD_600_; the strain was hypo-mucoid when evaluated by string test and sedimentation (**Fig 2B and S1A**).

### A reverse phenotypic screen mapped genes linked to capsular polysaccharide biosynthesis and hypermucoviscosity

A recent study used density-TraDISort to identify *K. pneumoniae* transposon mutants with altered buoyancy as a surrogate for CPS production and hmv [23]. This work identified transposon mutants that increased NTUH-K2044 buoyancy, and mutants that decrease NTUH-K2044 and/or ATCC 43816 buoyancy [23]. We sought to integrate the results of the forward screen (**Fig 1–2** and **Table 2**) with these results by systematically exploring the hmv and CPS production of KPPR1 transposon mutants in genes identified to have altered buoyancy. Nine of these genes were primary hits in the forward genetic screen and six passed the secondary and tertiary screens, including *pgi, ompR, kvrB (mprA), wzi, wza*, and *rmpA* (**Fig 1–3, Table 2**). Altogether, 56 mutants identified by density-TraDISort to impact *K. pneumoniae* buoyancy [23] were revived from the KPPR1 condensed library and evaluated for hmv by sedimentation and CPS production by uronic acid quantification (**Fig 3**). Twenty of these mutants had been identified to increase buoyancy in NTUH-K2044 and 36 of these mutants had been identified to decrease buoyancy in NTUH-K2044 and/or ATCC 43816 [23]. Note that KPPR1 is a rifampin-resistant derivative of ATCC 43816 [32].

**Fig 3.**
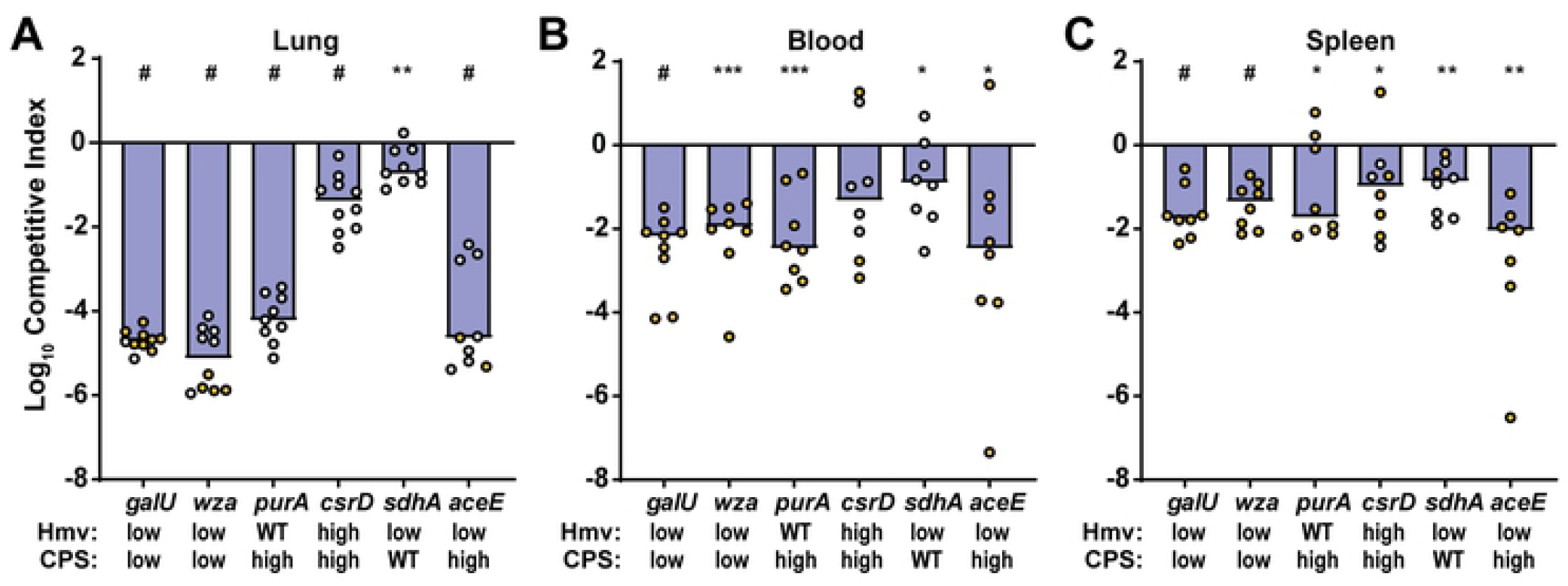
A reverse screen identifies mutants with altered mucoidy and capsule levels in KPPR1. Strains reported to have reduced or increased buoyancy were revived from the ordered KPPR1 transposon library. (A, C) The amount of capsule produced by each mutant was quantified by measuring uronic acid content and normalized to the OD_600_. (B, D) The percent of hypermucoid cells that remain in suspension were quantified after low-speed centrifugation. The x-axis labels in (B, D, F) apply to (A, C, E). The transposon insertion site for each mutant is labeled with the old locus tag number and gene name. Genes originally identified to decrease buoyancy in ATCC 43816 are navy, in NTUH-K2044 are light blue, or both are white; genes identified to increase buoyancy in NTUH-K2044 are yellow. All error bars represent one standard deviation from the mean and each assay was performed with three or more replicates. Statistical significance between wildtype (WT) (gray bar or line) and each mutant was determined using the Holm-Sidak method, with alpha = 0.05. Computations assumed that all rows were sampled from populations with the same scatter where, * P < 0.05; ** P < 0.01; *** P < 0.001; # P < 0.0001. The mean WT value is denoted by a horizontal, dotted gray line. The (E) capsule production and (F) hypermucovisity of each revived KPPR1 transposon mutant was classified according to how they were originally identified in [23]. Each circle represents an individual transposon mutant from A-D. Error bars represent one standard deviation from the mean and statistical differences between WT [in (E) N = 45 and in (F) N = 30] and classes of mutants were determined by unpaired *t*-test where, ** P < 0.01 and # P < 0.0001.

Of the 36 genes predicted to decrease CPS production, 14 had significantly reduced hmv and/or CPS. Six were significantly hmv^low^/CPS^low^ (*galU, rfaH, wzyE, arnD, arnE, wcaJ*), three were hmv^WT^/CPS^low^ (*rnfC2, arcB, pgm*), and five were hmv^low^/CPS^WT^ (*uvrY, miaA, galF, arnF, orf2*); surprisingly, one was hmv^WT^/CPS^high^ (*rnfD*) and two were hmv^high^/CPS^WT^ (*rnfE, gnd*) (**Fig 3A-B** and **Table 3**). Of the 20 genes previously identified to increase CPS production in NTUH-K2044, two were hmv^high^/CPS^high^ in KPPR1 (*pitA, csrD*), while four were hmv^WT^/CPS^high^ (*sapA, cyaA, polA, ptsN*) (**Fig 3C-D** and **Table 3**). Intriguingly, ten transposon mutants trended toward hmv^low^/CPS^high^ (*uvrY, rnfD, orf2, sapB, hha, aceE, purA, smpB, pta2, ackA3*) (**Fig 3A-D**). Overall, transposon insertions in genes previously determined to reduce buoyancy in both ATCC 43816 and NTUH-K2044, were most likely to reduce hmv or CPS in KPPR1 (8 of N = 12, 66.7% validation) [**Fig 3A-B** (white bars), **E-F** and **Table 3**]. This means that only six other genes (N = 24, 25.0%) previously identified to reduce buoyancy in ATCC 43816 or NTUH-K2044 validated with the KPPR1 transposon mutants present in the condensed library [**Fig 3A-B** (navy and light blue bars) and **Table 3**]. These results emphasize that genes identified across multiple strains are more likely to be integral to CPS biosynthesis and hmv biology species-wide. Furthermore, when evaluated as a whole group, genes previously identified to increase buoyancy significantly increased CPS levels, but not hmv, in KPPR1 transposon mutants (**Fig 3E-F**). In total, these results echo what was observed in the forward screen, that hmv and CPS overproduction are two distinguishable phenotypes (**Fig 2–3**).

**Table 3.**
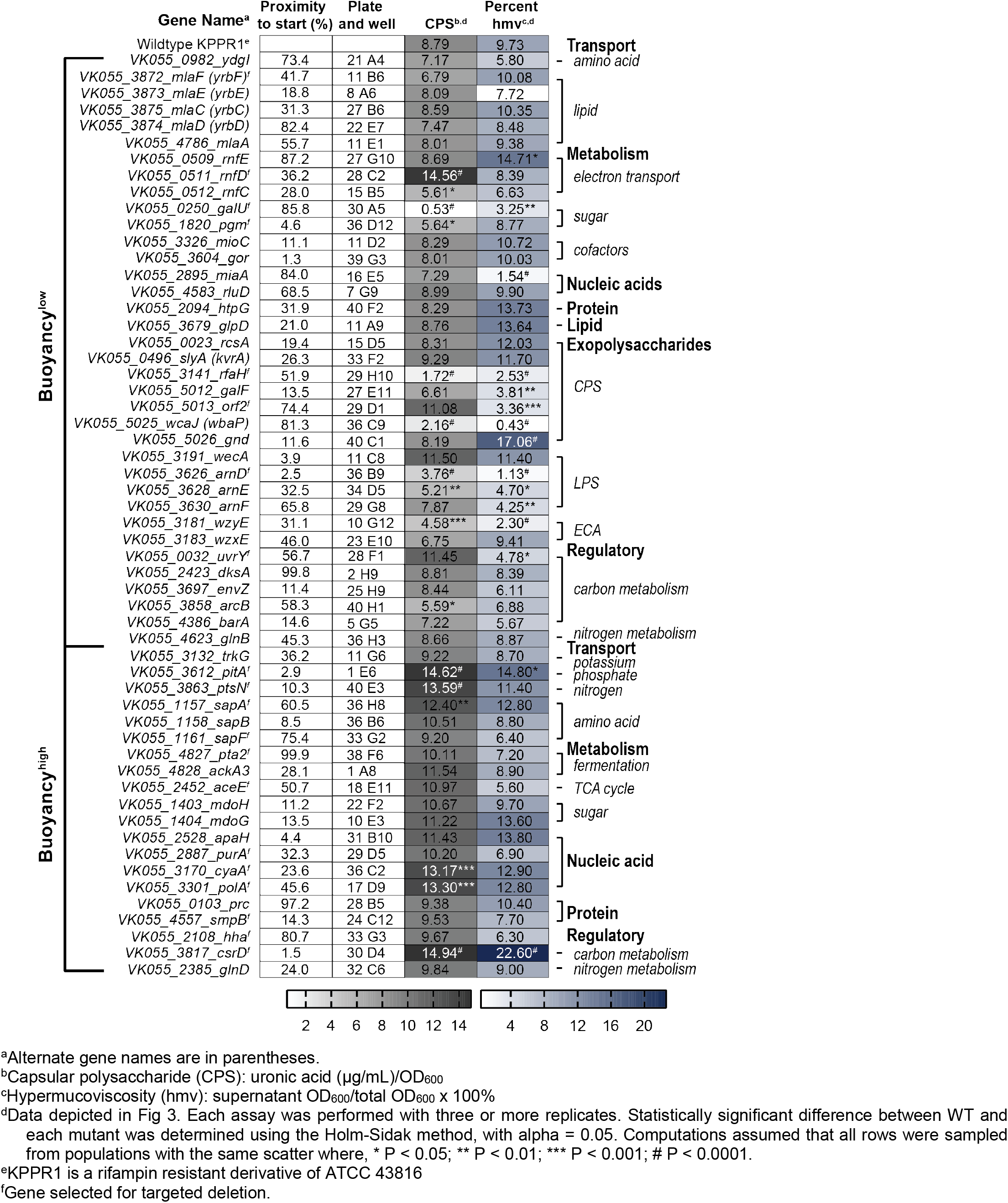
Reverse phenotypic screen of KPPR1 hypermucoviscosity and capsular polysaccharide biosynthesis.

### Hypermucoviscosity and CPS production are coordinated, but dissociable

All together, the forward and reverse screens quantified both hmv and CPS production in 100 transposon mutants and identified 43 hmv^low^/CPS^low^ mutants, three hmv^WT^/CPS^low^ mutants, five hmv^low^/CPS^WT^ mutants, two hmv^high^/CPS^WT^ mutants, one hmv^WT^/CPS^high^ mutant, and one hmv^WT^/CPS^high^ mutant. These data provide a rich resource for examining if *K. pneumoniae* CPS production and hmv are indeed interconnected. The nonparametric Spearman correlation coefficient between uronic acid concentration and sedimentation efficiency for all 100 transposon mutants examined was *r*^2^ = 0.5924 (p < 0.0001), identifying a significant link between the two variables.

To confirm that the phenotypes observed in the transposon mutants identified in the forward and reverse screens are attributable to the disrupted gene, a subset of these 100 transposon mutants were identified for further study. Twenty-seven representative transposon insertions were selected for targeted mutagenesis based on having diverse combinations of CPS levels and hmv (**Fig 2–3** and **Tables 2–3**). The resulting isogenic mutants were then systematically evaluated for CPS production and hmv (**Fig 4A-B**) [33]. Seventeen isogenic mutants (63.0%) exhibited significantly altered CPS production and hmv similar to the corresponding transposon mutant. These 17 isogenic mutants fell into six categories: (1) hmv^low^/CPS^low^ (Δ*uvrY*, Δ*galU*, Δ*rfaH*, Δ*VK055_3211*, Δ*arnD*, Δ*wza*), (2) hmv^WT^/CPS^low^ (Δ*pgm*), (3) hmv^WT^/CPS^high^ (Δ*hha*, Δ*aceE*, Δ*purA*, Δ*smpB*, Δ*pta2*), (4) hmv^low^/CPS^WT^ (Δ*sdhA*), (5) hmv^high^/CPS^high^ (Δ*cyaA*, Δ*polA*, Δ*csrD*), and (6) hmv^low^/CPS^high^ (Δ*aceE*) (**Fig 4**). The hmv and CPS quantification data from Fig 4A and 4B were aggregated on a single X-Y plot to evaluate the relationship between hmv and CPS production in the targeted mutants (**Fig 4C**). The nonparametric Spearman correlation coefficient between uronic acid concentration and sedimentation efficiency for all 27 targeted deletion mutants was *r*^2^ = 0.8041 (p < 0.0001), again supporting the historical perspective that CPS production and hmv are interconnected processes. However, it is notable that several mutants only had one parameter significantly change (Δ*sdhA*, Δ*purA*, Δ*pgm*, Δ*hha*, Δ*smpB*) or, surprisingly, had CPS production and hmv significantly altered in opposite directions (Δ*aceE*) (**Fig 4**). Moreover, we did not identify any mutants with increased hmv and reduced or WT levels of CPS biosynthesis, supporting the requirement of CPS biosynthesis for hmv.

**Fig 4.**
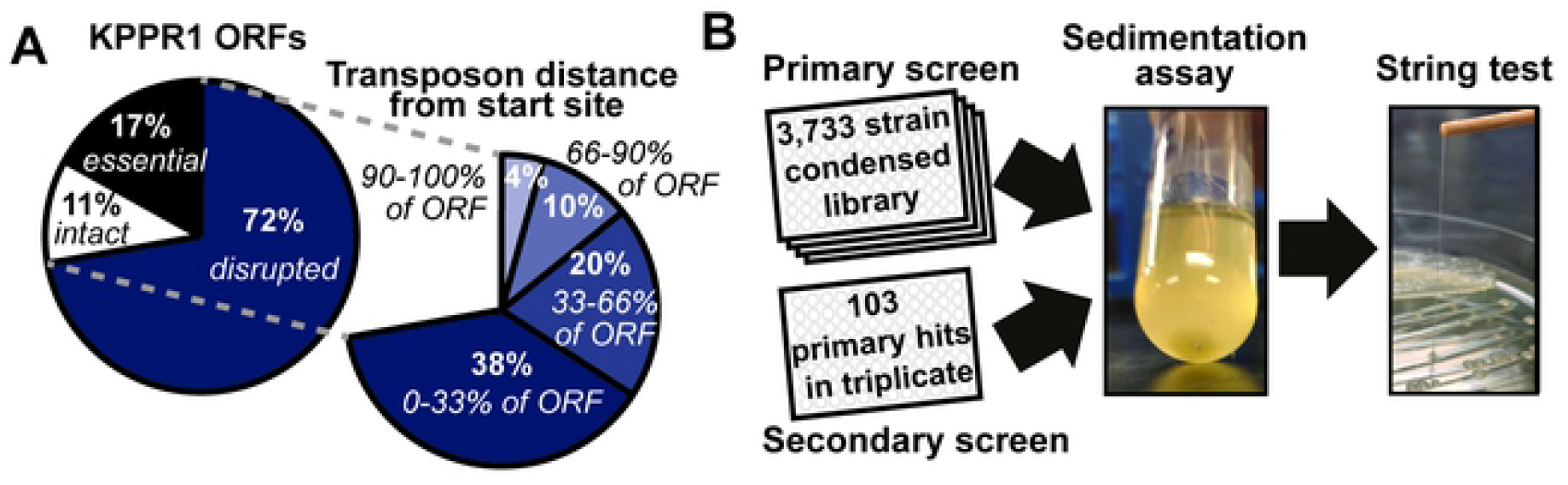
Hypermucoviscosity and capsular polysaccharide levels are coordinated, but dissociable. (A) The amount of capsule produced by 27 targeted deletion mutants was quantified by measuring uronic acid content and normalized to the OD_600_. (B) The percent of hypermucoid cells remaining in suspension were quantified after low-speed centrifugation of 1 OD_600_ unit of cells resuspended in 1 ml of PBS. All error bars represent one standard deviation from the mean and each assay was performed with six or more replicates. Statistical significance between wildtype (WT) and each mutant was determined using the Holm-Sidak method, with alpha = 0.05. (C) Data from A and B were coordinately plotted on a single graph and labeled with the gene name or gene number (VK055_XXXX). The nonparametric Spearman correlation coefficient for all targeted deletion mutants is r^2^ = 0.8041#. All computations assumed that data were sampled from populations with the same scatter where, * P < 0.05; ** P < 0.01; *** P < 0.001; # P < 0.0001. The mean WT value is denoted by dotted gray lines and a black marker.

Five mutants (Δ*galU*, Δ*wza*, Δ*purA*, Δ*csrD*, Δ*sdhA*), representing an array of altered CPS and hmv levels, were complemented *in trans* with the deleted gene under the control of its native promoter. For all mutants with altered CPS production, the complementation vector restored CPS production to WT levels, indicating that the altered CPS production of these strains is not due to secondary mutations in the chromosome (**Fig S2**). Although pACYC184 is a low copy number plasmid, complementing Δ*sdhA in trans* significantly reduced CPS levels compared to WT or Δ*sdhA* with empty vector (**Fig S2**). It is possible that CPS levels are very responsive to SdhA activity and the complementation vector does not fully recapitulate WT-level function.

### Isogenic mutants with altered CPS production and hypermucoviscosity are less fit in a murine pneumonia model

It is well-established that *K. pneumoniae* requires CPS to be fully virulent in multiple models of *K. pneumoniae* infection, including pneumonia and UTI [21, 22]. We hypothesized that both CPS production and hmv are important for full virulence and that disconnecting the two processes may reduce *in vivo* fitness. To test this hypothesis, six mutants encompassing a variety of hmv and CPS combinations, including hmv^low^/CPS^low^ (Δ*galU* and Δ*wza*), hmv^WT^/CPS^high^ (Δ*purA*), hmv^high^/CPS^high^ (Δ*csrD*), hmv^low^/CPS^WT^ (Δ*sdhA*), and hmv^low^/CPS^high^ (Δ*aceE*) were competed against WT KPPR1 in a murine model of disseminating pneumonia [33]. Mice were inoculated retropharyngeally with a targeted input ratio of 1:1 WT:mutant. At 24 h post-infection, bacterial burdens of WT and mutant in the lungs, blood and spleens were enumerated and the competitive indices were calculated (**Fig 5**). All mutants were significantly out-competed *in vivo*. Δ*galU* (53,500-fold), Δ*wza* (114,400-fold), Δ*purA* (15,200-fold), and Δ*aceE* (16,300-fold) were all dramatically out-competed in the lung (**Fig 5A**) and did not disseminate into the blood and spleens, as mutant CFUs were below the limit of detection in these organs (**Fig 5B-C**). Δ*csrD* (25.6-fold) and Δ*sdhA* (3.7-fold) had less dramatic decreases in competitive fitness in the lungs (**Fig 5A**) and were still able to disseminate into the blood and spleens of several mice (**Fig 5B-C**). Although all mutants were significantly outcompeted *in vivo*, they exhibited diverse growth phenotypes in LB medium (**Fig S3**). *In vitro*, only Δ*aceE* had a significantly longer doubling time than WT (3.5-fold) (**Fig S3A**). In addition, all strains except Δ*csrD* yielded less total bacterial growth than WT *in vitro*, as quantified by the area under the growth curve; although significant, these differences were quite subtle (**Fig S3B**). Altogether, these results suggest that appropriate coordination of CPS biosynthesis, hmv, and central metabolism is required for optimal fitness in a murine pneumonia model.

**Fig 5.**
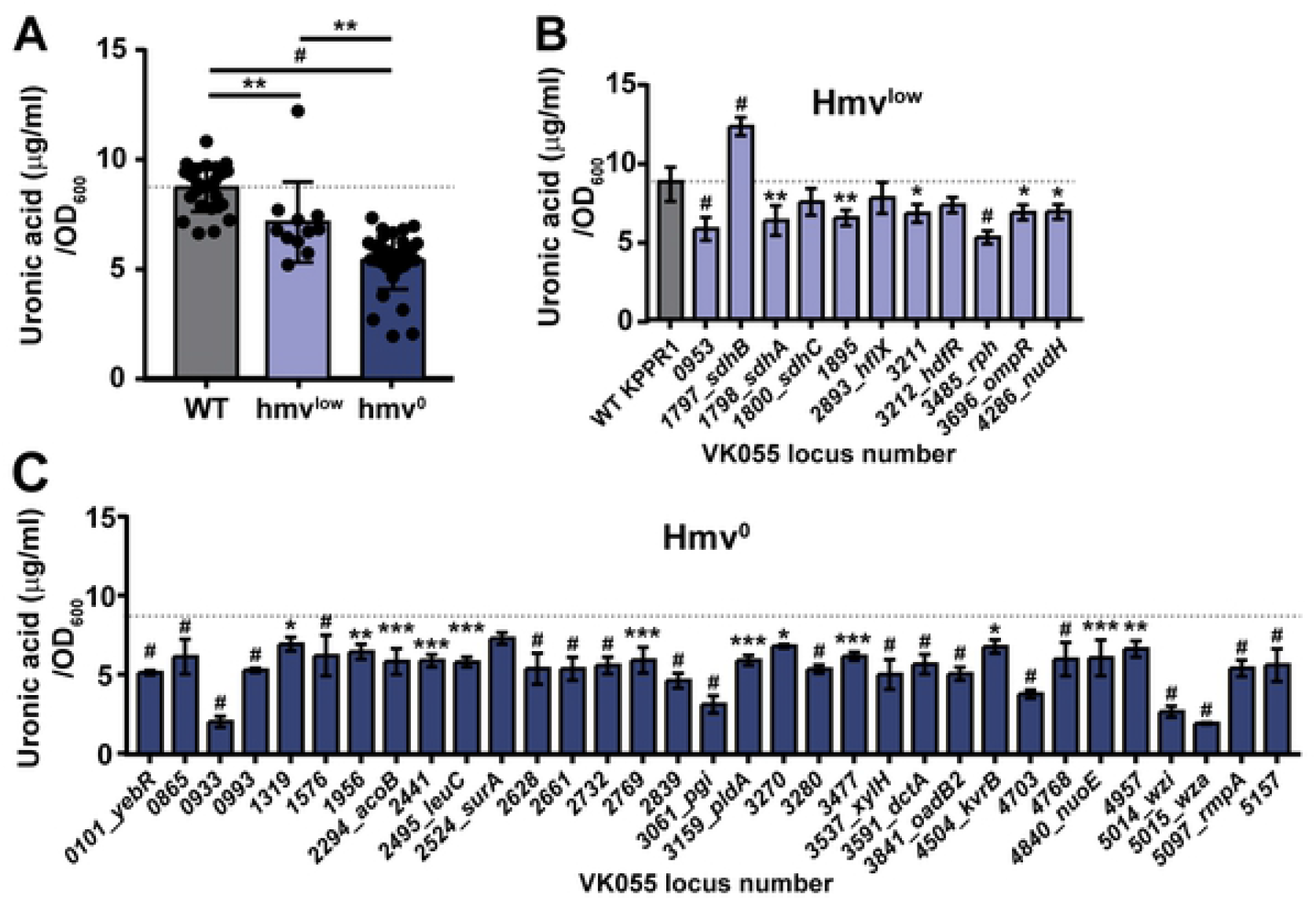
The *in vivo* fitness of select mutants is attenuated in a murine model of disseminating pneumonia. C57BL/6 mice were infected retropharyngeally with 1 x 10^6^ colony forming units (CFU) of a 1:1 ratio of wildtype (WT) to mutant, where each mutant and its significantly different levels of hypermucoviscosity (hmv) and capsular polysaccharide (CPS) are identified on the x-axis. The input ratios were determined by differential plating on LB and LB+kan. After 24 h of infection, the bacterial burdens of WT and each mutant in the (A) lungs, (B) blood, and (C) spleens were determined by differential plating. Each dot represents an individual mouse and yellow dots indicate that no mutant was detected in the outputs. The limit of detection was 100 CFU/ml. Competitive indices were calculated by dividing the output ratio of mutant/WT by the input ratio of mutant/WT. All competitive indices were log10-transformed and any significant differences from a competitive index of 0 was determined by a one-sample *t*-test where, * P < 0.05; ** P < 0.01; *** P < 0.001; # P < 0.0001.

## Discussion

CR-cKp tops the list of urgent antibiotic-resistant threats most recently released by the CDC [38]. Moreover, the growing incidence of hvKp and the emergence of CR-hvKp emphasizes the looming threat that *K. pneumoniae* poses to human health [3]. Historically, the hypervirulence of hvKp has been primarily attributed to RmpA-mediated increased production of CPS, along with stealth siderophores. These strains can often be identified by a positive string test, demonstrating hypermucoviscosity [3, 5]. The hmv of hvKp is generally ascribed to increased production of CPS; however, recent studies have challenged this paradigm [24–27, 30]. This paradigm shift in our understanding of *K. pneumoniae* hmv has been an emerging concept in recent years as genes, namely *kvrA, kvrB, rmpC*, and *rmpD*, have been identified that differentially affect hmv and CPS biosynthesis [17, 25, 26, 30]. Here, we have generated a condensed, ordered transposon library in a genetically tractable and hmv strain of *K. pneumoniae*, KPPR1, and used this library to query a question at the forefront of *K. pneumoniae* pathogenesis, namely, how do hmv and CPS biosynthesis independently and coordinately impact invasive hvKp infections?

In total, we quantified the impact of 100 transposon insertions and 27 targeted deletions on both CPS production and hmv. The relationship between these two properties was examined and the nonparametric Spearman correlation coefficient for CPS production and hmv was significant for both the transposon mutants (*r*^2^ = 0.5924; P < 0.0001) and targeted deletion strains (*r*^2^ = 0.8041; P < 0.0001). These data support the long-held view that hmv and CPS production are inter-related. However, several transposon and targeted deletion mutants dissociate CPS production and hmv, also supporting the emerging perspective that CPS is not the only biochemical feature driving hmv [17]. Considering these data altogether, we propose that in order to exhibit hmv, *K. pneumoniae* requires CPS along with other biochemical factors. Some targeted mutants with dissociated CPS production and hmv that may help dissect how these two processes intersect include pyruvate dehydrogenase, Δ*aceE* (hmv^low^/CPS^high^); succinate dehydrogenase, Δ*sdhA* (hmv^low^/CPS^WT^); and adenylosuccinate synthase, Δ*purA* (hmv^WT^/CPS^high^) (**Fig 4**). In addition, some transposon mutants exhibited phenotypes that trended toward increasing CPS production while reducing hmv (**Fig 2–3** and **Table 2–3**). Several genes related to the TCA cycle, pyruvate metabolism, and cellular energetics appear to decouple CPS biosynthesis and hmv. This suggests that hmv and CPS biosynthesis are integrated into the metabolic status of the cell. Further studies are required to dissect the metabolic pathways that control the biosynthesis of CPS and hypermucoviscosity. Moreover, the ability of the bacteria to distinctly control hmv and CPS biosynthesis suggest that there are environmental conditions in which one or both properties are advantageous, emphasizing that while these two features are closely associated with hypervirulent *K. pneumoniae*, they likely serve distinct functions within specific environments and may actually be regulated in response to the local environment [39]. For example, the hmv^low^/CPS^WT^ Δ*rmpD* mutant adheres to macrophage-like J774A.1 cells more than WT, while the hmv^WT^/CPS^low^ Δ*rmpC* mutant adheres similar to WT [25, 26]. In addition, non-mucoid strains are more likely to be isolated from urine than blood [39]. Hypothetically, hmv may be critical for evading adherence during invasive infections, but may interfere with critical adhesion factors during UTI.

To begin understanding the individual roles of hmv and CPS within the context of the host using the mutants generated here, we selected three targeted deletion mutants (Δ*aceE*, Δ*sdhA*, and Δ*purA*) with disproportionate CPS production and hmv, along with Δ*galU* and Δ*wza* (hmv^low^/CPS^low^) and Δ*csrD* (hmv^high^/CPS^high^), to evaluate *in vivo* fitness in a murine model of disseminating pneumonia. In the murine pneumonia model, the major fitness defect and inability to disseminate from the lung to the blood and spleen for Δ*galU* and Δ*wza* supports the established importance of hmv and capsule *in vivo*, especially for invasive infections (**Fig 5**) [39–42]. The remaining four mutants, Δ*csrD*, Δ*aceE*, Δ*sdhA*, and Δ*purA*, have more complex biology. In addition to exhibiting altered hmv and/or CPS biosynthesis, all are involved in central metabolism, confounding any definite conclusions about how their protein products contribute to *in vivo* fitness. In fact, other purine biosynthesis mutants (*purF, purL, purH*) have been identified to have a fitness defect in *K. pneumoniae*; although, the impact of these mutations on CPS production and hmv was not evaluated [33]. The predicted alterations in carbon metabolism and cellular redox status in these mutants may itself alter the *in vivo* fitness of *K. pneumoniae*, but it is also possible that the observed fitness defects are due to altered CPS biosynthesis and hmv. It is intriguing that the two mutants (Δ*csrD* and Δ*sdhA*) that retain the ability to disseminate from the lungs are those that maintain ratios of CPS to hmv most similar to WT. This observation suggests that both CPS production and hmv may be important for invasive *K. pneumoniae* infections, which has been observed clinically [39]. Nonetheless, further studies are needed to identify the precise signals that regulate CPS biosynthesis and hmv, as well as dissect how these regulatory pathways overlap, diverge, and impact pathogenesis.

Some novel pathways identified here that should be of immediate focus are the succinate dehydrogenase complex and the hypothetical gene *VK055_3211* and its divergently transcribed regulator, *VK055_3212_hdfR* [43, 44]. Homologues to *VK055_3211* and *hdfR* have been identified in *E. coli* to contribute to the organization of the Ori region during chromosome replication and HdfR has been shown to repress the flagellar master operon (*flhDC*) [43, 45]. Although KPPR1 does not encode *flhDC, hdfR* expression is repressed by H-NS, which has been shown to repress hmv and CPS biosynthesis in *K. pneumoniae* [45, 46]. Altogether, these data suggest that HdfR may be another component of the complex CPS biosynthesis and hmv regulatory networks in *K. pneumoniae* and may coordinate these features with cell replication. More globally, the identification of genes linked to central metabolism that, when disrupted, result in a decrease in hmv suggests that hmv is tightly linked to the energy status of the cell. This is not too surprising as elaborating large extracellular macromolecules is an energetically expensive process and has been showed to serve as an energy reservoir in other bacterial species [47–49]. It is intriguing that many of these hits result in complete loss of hmv, while retaining intermediate levels of CPS production (**Fig 2**). The stronger effect of perturbing bioenergetics on hmv than CPS may explain why these genes have not previously been identified to impact CPS biosynthesis.

Further strengthening the connection between the integration of cellular metabolism with the regulation of CPS and hmv, is our confirmation that several genes involved in the carbon storage regulatory network coordinately increase buoyancy, where BarA/UvrY and DksA oppose CsrD and CyaA activity [23, 50]. By systematically quantifying hmv and CPS production in transposon mutants previously identified to impact buoyancy in NTUH-K2044 [23], we have confirmed that transposon insertions in *uvrY* significantly decreases hmv, *cyaA* increases CPS, and *csrD* increases both CPS biosynthesis and hmv in KPPR1 (**Fig 3A-D**). The carbon storage regulatory network interfaces with the cAMP receptor protein, CRP, which has been previously shown to repress CPS biosynthesis at the transcriptional level in NTUH-K2044 and CG43 [51, 52]. On the other hand, of all the tested genes in the *sapABCDF* cationic peptide ABC transporter operon, which had been identified to increase buoyancy in NTUH-K2044, only *sapA* significantly increased CPS levels in KPPR1 (**Fig 3A-D**). Altogether these results suggest that the carbon storage regulatory circuit may represent a more broadly conserved mechanism *K. pneumoniae* employ to control hmv and CPS biosynthesis, while the *sap* operon may exert control of these processes in clonal groups more closely related to NTUH-K2044. It is important to appreciate that while the density-TraDISort study was only able to identify transposon mutants that increased CPS production in NTUH-K2044, many had a similar effect in KPPR1. The authors did note that several genes including *uvrY, barA, csrB, rcsA* and *rcsB* met some, but not all, of their screening criteria to be identified as hits in ATCC 43816 [23]. Altogether, the results of the forward and reverse screens further support the notion that CPS biosynthesis and hmv are tightly linked to the metabolic state of *K. pneumoniae* and that although hmv requires CPS production, it is not the only factor. Therefore, it is critical to continue to evaluate both of these virulence-associated features so that biological effects on each process may be assessed independently. This may be accomplished by focusing on hits identified here that only affect hmv or CPS biosynthesis, or in some cases impose an opposite effect on these two properties. It may be that changes in the intracellular pools of metabolic intermediates or signaling nucleotides in response to environmental oxygen, carbon- or nitrogen-sources differentially regulate hmv and CPS.

The reverse screen executed here built on a recent density-TraDISort study that identified transposon mutants with altered buoyancy in NTUH-K2044 and/or ATCC 43816, the parental strain of KPPR1 [23]. For those transposon mutants that did not reproduce the previously reported phenotype, it is important to appreciate that the two screens are experimentally distinct in that one was performed by separating a pool of mutants over a Percoll gradient and the other probed each mutant individually using the sedimentation assay. Some mutants may conceivably behave differently when assayed in a pool versus individually. This is especially true if the product of the mutated gene can be complemented by other mutants in the pool that are effectively WT for the gene of interest. Alternatively, it is possible that the transposon mutants in the KPPR1 library are not relevant under the experimental conditions or functionally inactivating. However, the site of transposon insertion ranged from 1.3-99.8% from the predicted start codon, with a median value of 34.35% (17.75-66.48% interquartile range). Thus, most transposon mutants are expected to be functionally disrupted (**Table 3**). For those genes that did not exhibit an effect on CPS or hmv, the median distance from the start site was 31.3% (17.75-45.825% interquartile range), indicating that most negative results skewed toward the start codon. This observation was surprising as we anticipated that transposon insertions toward the end of the gene, would be more likely to have less impact on function. This expectation likely over-simplifies the complexities of protein function and operon structure and suggests that many of the transposon mutants in the condensed library provide valuable biological insights, regardless of their distance from the start site. It may even be valuable to return to the full transposon library to access multiple transposon insertion sites in the same gene or operon, thereby providing a comparison of similar, but unique mutants. Even fuller datasets may be achieved by accessing the two other ordered *K. pneumoniae* transposon libraries, in addition to the one generated here. One such library contains 12,000 strains that correspond to 4,583 ORFs in strain KPN1H1 with the KPC-3 carbapenemase gene deleted and the other library has approximately 4,570 mapped transposon insertion site in ATCC 43816, although the number of unique ORFs disrupted is unclear [53, 54]. Altogether, these libraries represent invaluable genetic resources that will not only advance our understanding of *K. pneumoniae* pathogenesis and molecular biology within these specific strains, but can also be used as templates to generate insertional mutants in other strains or to study the contribution of individual domains to phenotypes of interest. Furthermore, a small, condensed library, as described here, provides an invaluable tool for circumventing bottle necks during *in vivo* TnSeq studies.

In summary, we have generated a rich data set of mutants with a range of effects on CPS biosynthesis and hmv. These data provide a framework for future studies focused on identifying the precise signals that regulate CPS biosynthesis and hmv, as well as dissecting how these two major features of hvKp independently and coordinately impact pathogenesis. We have shown that CPS biosynthesis and hmv are coordinated processes that can be dissociated by deleting genes tied to central metabolism. The assembly of CPS and formation of hmv are energetically expensive processes, so it is intuitive that these processes are hardwired to the metabolic pulse of the cell. The linkage between hypervirulent and invasive *K. pneumoniae* and its overproduction of CPS and hmv may provide a fitness advantage for invasive infections, but at a metabolic cost. It is possible that in more stringent environments the metabolic burdens of elevated CPS biosynthesis and hypermucoviscosity may pose a fitness disadvantage. This cost-benefit balance between adequate energy sources and resisting environmental stresses, such as a healthy immune response, may explain the emergence of the hvKp lineage and its invasive pathology compared to cKp strains.

## Materials and Methods

### Bacterial strains and media

*Klebsiella pneumoniae* strain KPPR1, a rifampin-resistant derivative of ATCC 43816, was used for all studies [32]. All primers, strains and plasmids described in these studies are detailed in **Supplemental Tables 1** and **2.** Bacteria were cultured in lysogeny broth (LB) (5 g/L yeast extract, 10 g/L tryptone, 0.5 g/L NaCl) at 200 rpm and 37 °C, unless otherwise noted. When appropriate, antibiotics were added at the following concentrations, rifampin (30 μg/mL), kanamycin (25 μg/mL), chloramphenicol (80 μg/mL), and spectinomycin (50 μg/mL). *Escherichia coli* strain TOP10 was used to generate complementation vectors and cultured in LB supplemented with chloramphenicol (20 μg/mL).

### Transposon library construction and sequencing

A library of random transposon mutants was generated in *K. pneumoniae* KPPR1 by conjugation with *E. coli* S17 harboring pSAM_Cam with a modified Mariner *Himar1* transposon as previously described [33]. Briefly, mid-log cultures of the donor and recipient strains were mixed in a 2:1 ratio, washed with PBS, resuspended in LB medium, and spread on filter disks on top of an LB agar plate. Following a 2 hr incubation at 37 °C, filters were transferred to an agar plate containing 250 μM IPTG (Invitrogen, Carlsbad, CA) and incubated for 2.5 hr at 37 °C to induce expression of the transposase, enabling mobilization of the transposon. Bacteria were resuspended in LB medium transferred from the filter to LB agar with rifampin (30 μg/mL) and kanamycin (50 μg/mL) to select KPPR1 isolates with genomic transposon insertions. Rifampin-, kanamycin-resistant trans-conjugants were inoculated into 192, 96-well microplates containing 200 μL LB medium with 15% (v/v) glycerol and 50 μg/mL kanamycin and incubated statically at 37 °C until saturation.

To verify that rifampin-, kanamycin-resistant colonies did not result from integration of the conjugation plasmid pSAM_Cam, a subset of the library was subjected to colony PCR utilizing primer pairs with homology to the plasmid backbone and the transposon as described previously (N = 10) [33]. The library was also tested to ensure that each mutant contained a single transposon insertion and that the insertion location was random by subjecting EcoRI-digested genomic DNA to Southern blotting using a probe homologous to the transposon as described previously (n = 13) [33].

Identification of the location of the transposon insertion site within the KPPR1 chromosome for each individual mutant was accomplished using next-generation sequencing coupled with Cartesian pooling to reduce the total number of samples to be sequenced. Using the method presented in [31], the number of samples to be sequenced is condensed first from 18,432 wells to 80 mutant pools, representing the physical location of the mutants in the X, Y, and Z planes within each stack of 96, 96-well microplates. Each condensed pool of mutants is assigned a 6 bp barcode and the representation of each mutant within a barcoded pool is used to de-convolute the physical location within the library. To generate the mutant pools, transposon library plates were replicated into 75 μL of LB + 25 ug/mL kanamycin (LB+kan) medium and cultured statically at 37 °C overnight. The following day, 75 μL of 50% sterile glycerol was added to each plate, then Cartesian pooling was executed as previously described [31]. All intermediate mutant pools were stored at −20 °C. Intermediate XY and Z pools were thawed and combined [31].

Genomic DNA was isolated from 1 mL of the combined final XY and Z pools using the DNeasy Blood and Tissue kit according to the manufacturer’s directions for gram-negative bacteria (Qiagen). Genomic DNA (1 μg) for each mutant pool was sheared using a Covaris DNA fragmentation system (Intensity = 5; duty cycle = 5%; cycles per burst = 200; 55 s), resulting in an average fragment size of 370 bp and ranging from 200-700 bp. Sheared DNA was blunt-end repaired and dA-Tailing was added using the NEBNext Ultra-End Repair/dA-Tailing Module. The DNA was then purified using AMPure XP, eluting in 25 μl water. All down-stream library preparation and sequencing data analysis was performed as described previously (**Supplemental Table 1**) [31, 55].

### Condensed library construction

The KPPR1 genome (GCA_000742755.1) was used to identify predicted ORFs and gene coordinates and the fgenesB predictor was used to identify predicted transcriptional units [56]. The TP ID for each transposon insertion identified was manually matched to the plate number as described in [31]. The Fuzzy Join function of the Fuzzy Lookup Add-In for Microsoft Excel was then used to match the transposon insertion sites, plate (TP) and well (AP) coordinates, gene name, and gene coordinates. All Fuzzy Join functions had a similarity threshold = 1. The percent of the gene disrupted by the transposon insertion was calculated using the following two equations, where **Eq. 1** was used for genes on the positive-strand and **Eq. 2** was used for genes on the negative-strand:

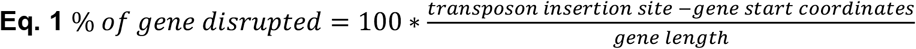

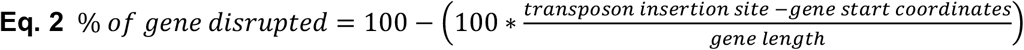

Plate and well coordinates with multiple mutants mapped to the location were identified and counted using basic Excel functions.

The ordered library was then curated to identify optimal transposon mutants to be included in the condensed, ordered library. All intergenic mutations were removed from the data set and the resulting data set was sorted by gene name and then by percent of the gene disrupted. Fuzzy Join was then used to identify 1 transposon mutant for each gene in the KPPR1 genome. This condensed library was then evaluated and hand-curated to ensure that the positional location of the transposon mutant selected for the condensed library was identified with high confidence and contained a single insertion site, if possible. Selected transposon mutants were re-arrayed into microplates containing LB+kan medium, grown statically overnight at 37 °C, mixed with an equal volume of 50% glycerol then stored at −80 °C to make the condensed, ordered library.

### Hypermucoviscosity sedimentation assays

The hmv was assessed as described previously with the following modifications [33]. The overnight cultures were pelleted at 21,000 *x g* for 15 min then resuspended to an OD_600_ = 1.0 in a final volume of 1 mL PBS. Samples were centrifuged at 1,000 *x g* for 5 min and the OD600 of the upper 900 μL supernatant was determined in a 1 cm cuvette.

### Uronic acid quantification

Analysis of the total uronic acid content was performed following a modified procedure [57]. A 0.25 mL volume of overnight culture was mixed with 50 μL 1% Zwittergent 3-14 in 100 mM citric acid buffer, pH 2 at 50 °C for 20 min. Bacterial cells were pelleted by centrifugation then 0.1 mL of the cell-free supernatant was mixed with 0.4 mL absolute ethanol and incubated according to [57]. Samples were rehydrated in 0.2 mL of water then 1.2 mL of 0.0125 M sodium tetraborate in concentrated sulfuric acid was added. All subsequent steps were as described in [57] and normalized to the total OD_600_.

### Forward screen

Microplates containing the condensed, ordered library (total of 3,733 mutants) were thawed at room temperature and replicated into 100 μL of LB in round bottom microplates. Plates were wrapped with plastic wrap to prevent evaporation and incubated statically at 37 °C for 18-19 h. The sedimentation assay was adapted to a microplate format as follows. Plates were vortexed on low for 60 sec then the total OD_600_ was recorded. Plates were centrifuged at 2,000 *x g* for 20 min, then the upper 50 μL of supernatant was transferred to a new microplate to measure the OD_600_. Transposon mutants with a total OD_600_ less than two standard deviations from the plate mean and a supernatant OD_600_ more than two standard deviations from the plate mean were considered hits. The hits were struck onto LB agar, incubated at 37 °C overnight and evaluated by string test the following day. Three colonies of each transposon mutant confirmed as non-mucoid or hypo-mucoid by string test were arrayed into a microplate for confirmation. The same work-flow with sedimentation and string test were repeated with the arrayed hits for confirmation. The top hits were confirmed in a third sedimentation assay where the transposon mutants were cultured in 3 mL of LB medium overnight at 37 °C with aeration, then the OD_600_ of 100 μL of the total culture and culture supernatant, after 7,000 *x g* for 10 min, was determined. Pathway analysis was performed using KEGG GENES (Kyoto Encyclopedia of Genes and Genomes) [36, 37].

### Reverse screen

Transposon mutants were revived on LB agar plates, then individual colonies were inoculated into 3 mL of LB medium and incubated overnight at 37 °C with aeration. Uronic acid quantification was performed as described above in parallel with a modified sedimentation assay. The sedimentation assay was performed by recording the OD_600_ of 100 μL of overnight culture in a microplate, followed by pelleting 1 mL of the overnight culture at 7,000 *x g* for 10 min, and quantifying the OD_600_ of the upper 100 μL of the culture. The ratio of supernatant to total OD_600_ was used as a measure of hmv.

### Construction and complementation of mutants

Insertional mutants were generated using λ Red recombineering adapted to *K. pneumoniae* as described previously with the following exceptions [33, 58]. All bacterial cultures for competent cells were supplemented with 0.5 μM EDTA, which improves centrifugation. Electrocompetent cells were either transformed immediately or flash frozen and stored at −80 °C for future use. PCR products with 60 base pairs of homology flanking the region targeted for deletion were digested with DpnI and 6 μL of the column purified PCR product was mixed with competent KPPR1 pKD46 cells and incubated on ice for 30 min. Transformants were recovered with 500 μL of LB and static incubation at room temperature overnight, although some mutants required recovery at 30 °C for 3-4 hr or 37 °C for 1-2 h, with shaking.

All mutants were generated using pKD4 template, which confers kanamycin resistance. Successful mutagenesis was confirmed by PCR and restriction digest with EagI. All oligonucleotides for mutagenesis and confirmation are listed in **Supplemental Table 1**.

Complementation vectors were generated using NEBuilder HiFi DNA Assembly Cloning Kit (New England BioLabs). Primers were designed using the online NEBuilder Assembly Tool with the following setting: >20 nucleotide overlap, Phusion DNA Polymerase (HF Buffer), 500 nM primer concentration (**Supplemental Table 1**). The ORF fused to 500 bp of the predicted promoter region were exchanged with 600 bp of the *tet* cassette in pACYC184 [59]. Gel purified PCR products were assembled according to the manufacturer’s instructions and the enzymatic reaction was incubated at 50 °C for 1 h. The NEBuilder reaction was dialyzed overnight against 10% sterile glycerol using a VSWF 0.025 μm filter disk. The dialyzed DNA was collected and electroporated into *E. coli* TOP10 cells. pACYC184 empty vector was generated by ligating the pACYC184 PCR product without an insert, effectively eliminating 600 bp of the *tet* cassette. The resulting plasmids were verified by restriction digest and Sanger sequencing then 0.5 μL of DNA was transformed into 50 μL of electrocompetent *K. pneumoniae* mutants [33].

### Growth analyses

Bacterial strains were cultured statically overnight in triplicate in 100 μL of LB medium in a microplate at 37 °C. The cultures were normalized to an OD_600_ of 0.01 in LB medium then 100 μL was aliquoted into a microplate. A Bioscreen-C Automated Growth Curve Analysis System (Growth Curves USA) was used to record the OD_600_ every 15 min for 24 h. Cultures were incubated at 37 °C with continuous, medium shaking. The doubling time was determined by identifying two time points (t2 and OD2 = late time point and t1 and OD1 = early time point) within the logarithmic growth phase, then applying **Equation 3**:

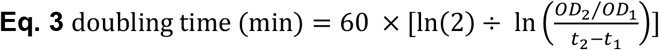

### Murine pneumonia model

A murine model of *K. pneumoniae* infection was used as previously described [33]. Briefly, 6-8 week/old C57BL/6 mice (Jackson Laboratory, Bar Harbor, ME) were anaesthetized with isoflurane and retropharyngeally inoculated with 1 x 10^6^ CFU *K. pneumoniae* in 50 μL of PBS. All bacterial strains were cultured overnight in 50 mL LB, except *wza::kan* was cultured in LB+kan. Bacteria were pelleted at 10,000 x *g* for 30 min and the pellets resuspended in sterile PBS to a final OD_600_ of 2.0. WT and mutant were mixed at a 1:1 ratio and the input colony forming units (CFU) ratios determined by serial dilution and drip plating on both LB and LB+kan. Infections were allowed to proceed for 24 hr and mice were euthanized by CO_2_ asphyxiation. Blood was collected by cardiac puncture in heparinized tubes. Lungs and spleens were collected and homogenized in 3 mL of sterile PBS. Whole blood and homogenized lungs and spleens were serial diluted in PBS and 10 μL drip plated on LB and LB+kan. Plates were incubated at 30 °C overnight and the CFUs enumerated the following morning. The limit of detection was 100 CFU/mL and all samples without detectable CFU counts were analyzed assuming that they contained 99 CFU/mL. The competitive index (CI) was calculated as in **Eq. 4**.

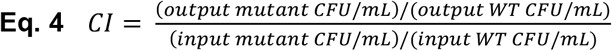

### Statistical analysis

All *in vitro* replicates represent biological replicates and all *in vivo* studies were replicated at least twice. All statistical analyses were computed in Prism 8.3.0 (GraphPad Software, La Jolla, CA). For *in vitro* experiments, significance was calculated using unpaired *t*-tests using the Holm-Sidak method to correct for multiple comparisons with alpha = 0.05. A two-tailed P value for the correlation between hmv and CPS production was computed by nonparametric Spearman correlation with a 95% confidence interval. For *in vivo* experiments, all competitive indices were log_10_ transformed then significance was calculated using a one sample *t-*test, where the actual mean was compared to a theoretical mean of 0.00 (no fitness defect). Results were considered significant if the P value was less than or equal to 0.05.

### Ethics statement

All animal studies were conducted in accordance with the recommendations in the *Guide for the Care and Use of Laboratory Animals* [60]. The University of Michigan Institutional Animal Care and Use Committee (IACUC) approved this research (PRO00007474).

## Author contributions

**LAM:** Conceptualization, Investigation, Data Curation, Formal Analysis, Methodology, Project Administration, Visualization, Writing – Original Draft Preparation, Writing – Review & Editing

**AJS:** Investigation, Data Curation, Formal Analysis, Methodology, Visualization, Writing – Review & Editing

**VF:** Conceptualization, Investigation, Data Curation, Formal Analysis, Methodology, Writing – Review & Editing

**JV:** Investigation, Methodology, Resources, Visualization, Writing – Review & Editing

**SNS:** Investigation, Writing – Review & Editing

**MAB:** Conceptualization, Funding Acquisition, Resources, Supervision, Writing – Review & Editing

**HLTM:** Conceptualization, Funding Acquisition, Project Administration, Resources, Supervision, Writing – Original Draft Preparation, Writing – Review & Editing

## Acknowledgments

This research was supported by Public Health Service grant R01 AI134731 from the NIH. We thank Alexandra Johnson for assistance with arraying transposon mutants; Juan Marzoa for technical advice and shared reagents; Caitlyn Holmes, Sébastien Crépin, and Allyson Shea for critical reading of the manuscript; and the Mobley and Bachman labs for critical feedback. We acknowledge support from the Advanced Genomics and Bioinformatics Cores of the University of Michigan Medical School’s Biomedical Research Core Facilities, specifically Dr. Weisheng Wu, who transformed the raw sequencing data into insertion site locations using the Galaxy platform and published code.

## Supporting Information Captions

**Fig S1. The centrifugation assay recapitulates the string test results of forward screen hits.** Hits in the forward screen were categorized as (A) hypo-mucoid (hmv^low^) or (B) non-mucoid (hmv^0^) by string test and sedimentation assays performed in microplates. To confirm that categorizing mutants based on these high-throughput methods is reflected in full-scale centrifugation assays, 3 mL of each mutant was grown overnight and centrifuged at 7,000 *x g* for 10 min. The optical density at 600 nm (OD_600_) of the supernatant was normalized to the total OD_600_ of the overnight culture by measuring the absorbance of 100 μL in a plate reader. Error bars represent one standard deviation from the mean of the assay performed in triplicate. Statistical significance between wildtype (WT) and each mutant was determined using the Holm-Sidak method, with alpha = 0.05. Computations assumed that all rows were sampled from populations with the same scatter. No results were significantly different from WT.

**Fig S2. Key mutants are complemented *in trans*.** Select targeted deletion mutants were transformed with either empty vector (pACYC184) or a complementation vector, which contained the targeted gene under the control of its native promoter on pACYC184. The amount of capsule produced by the empty vector and complementation strains was quantified by measuring uronic acid content and normalized to the OD600. All error bars represent the standard error of the mean and each assay was performed with nine or more replicates. Statistical significance between wildtype (WT) and each mutant was determined using the Holm-Sidak method, with alpha = 0.05. All computations assumed that data were sampled from populations with the same scatter where, # P < 0.0001. The mean WT empty vector value is denoted by a dotted gray line.

**Fig S3. *In vitro* growth of key mutants.** The *in vitro* growth of the six mutants co-inoculated with wildtype (WT) in a murine model of pneumonia was evaluated. Mutants were grown in LB and growth quantified by (A) measuring the optical density at 600 nm (OD_600_) each hour and (B) integrating the area under the growth curve in GraphPad Prism 8.3.0. Error bars represent the standard error of the mean and each data point represents at least nine replicates. Statistical significance between the area under the curve and doubling time of wildtype (WT) and each mutant was determined using the Holm-Sidak method, with alpha = 0.05. Computations assumed that all data points were sampled from populations with the same scatter where, * P < 0.05; ** P < 0.01; *** P < 0.001; # P < 0.0001.

**Table S1. Primers used in this study.**

**Table S2. Strains and plasmids used in this study.**

^a^Km = kanamycin; Cm = chloramphenicol; Tc = tetracycline; Rif = rifampin; Sp = spectinomycin

**Data Set S1. Transposon insertion sites and positional locations within the full and condensed ordered libraries.** (Tab 1) Legend for this data set. The nucleotide and gene location for each transposon insertion is reported in conjunction with the positional location and confidence with which that location was mapped for the full library. Transposons are reported based on if they (Tab 2) disrupt a single gene, (Tab 3) are intergenic, and (Tab 4) disrupt two genes. (Tab 5) All remaining genes not disrupted in the full library. (Tab 6) Map decoding the TP ID with the plate location for the full, ordered library. (Tab 7) Transposon mutants and their positional locations in the condensed library. All genes reported in this study are annotated using the old locus tags. (Tab 8) The old locus tag, its nucleotide location, and gene function have been matched with the new locus tags.

